# Physiological and molecular responses of a newly evolved auxotroph of Chlamydomonas to B_12_ deprivation

**DOI:** 10.1101/836635

**Authors:** Freddy Bunbury, Katherine E Helliwell, Payam Mehrshahi, Matthew P Davey, Deborah Salmon, Andre Holzer, Nicholas Smirnoff, Alison G Smith

**Author notes:** Corresponding author, Address: Department of Plant Sciences, Downing Street, Cambridge, CB2 3EA, UK.

## Abstract

The corrinoid B_12_ is synthesised only by prokaryotes yet is widely required by eukaryotes as an enzyme cofactor. Microalgae have evolved B_12_ dependence on multiple occasions and we previously demonstrated that experimental evolution of the non-requiring alga *Chlamydomonas reinhardtii* in media supplemented with B_12_ generated a B_12_-dependent mutant (hereafter metE7). This clone provides a unique opportunity to study the physiology of a nascent B_12_ auxotroph. Our analyses demonstrate that B_12_ deprivation of metE7 disrupted C1 metabolism, caused an accumulation of starch and triacylglycerides and a decrease in photosynthetic pigments, proteins and free amino acids. B_12_ deprivation also caused a substantial increase in reactive oxygen species (ROS), which preceded rapid cell death. Surprisingly, survival could be improved without compromising growth by simultaneously depriving the cells of nitrogen, suggesting a type of cross protection. Significantly, we found further improvements in survival under B_12_ limitation and an increase in B_12_ use-efficiency after metE7 underwent a further period of experimental evolution, this time in coculture with a B_12_-producing bacterium. Therefore, although an early B_12_-dependent alga would likely be poorly adapted to B_12_ deprivation, association with B_12_-producers can ensure long-term survival whilst also providing the environment to evolve mechanisms to better tolerate B_12_ limitation.

## Introduction

Over 50% of algal species require an exogenous source of B_12_ for growth (1), yet large areas of the ocean are depleted of this vitamin (2, 3). Eukaryotic algae cannot synthesise B_12_, but must instead obtain it from certain prokaryotes that can (1). Indeed, whilst dissolved B_12_ concentrations are positively correlated with bacterioplankton density (4, 5), they have been found to negatively correlate with phytoplankton abundance (6, 7). Furthermore, nutrient amendment experiments suggests B_12_ limits phytoplankton growth in many aquatic ecosystems (8–10). Despite this, understanding of the physiological and metabolic adaptations that B_12_-dependent algae employ to cope with B_12_ deprivation is rather limited.

In many algae B_12_ is required as a cofactor for the B_12_-dependent methionine synthase enzyme (METH) (11), although some algae encode a B_12_-independent isoform of this enzyme (METE) and do not require B_12_ for growth. Bertrand et al (12), showed that the B_12_-dependent marine diatom *Thalassiosira pseudonana*, which encodes only METH, responds to B_12_ scarcity by increasing uptake capacity and altering the expression of enzymes involved in C1 metabolism. Heal et al (13) found that despite these responses B_12_ deprivation disrupted the central methionine cycle, transulfuration pathway and polyamine biosynthesis. *Phaeodactylum tricornutum*, a marine diatom which uses but does not depend on B_12_ (encoding both METE and METH), responds similarly to *T. pseudonana* (12) but can also rely on increasing expression of METE to maintain the production of methionine. Phylogenetic analysis of the *METE* gene among diatoms shows no simple pattern of gene loss or gain, as indeed is the case across the eukaryotes (14, 15), but there is a clear link between the lack of a functional copy of the *METE* gene and B_12_-dependence (11, 16).

As with the diatoms, the phylogenetic distribution of *METE* within the *Volvocales* (a family of green freshwater algae) points to gene loss on several independent occasions. The genomes of two volvocalean algae, *V. carteri* and *G. pectorale*, contain *METE* pseudogenes indicating that B_12_ dependence has evolved relatively recently in these species (11). *Chlamydomonas reinhardtii* is a related alga that encodes a functional copy of *METE* and so is B_12_-independent. Helliwell et al. (17) generated a *METE* mutant of *C. reinhardtii* by experimental evolution in conditions of high vitamin B_12_ concentration, demonstrating that sustained levels of B_12_ in the environment can drive *METE* gene loss. This mutant, which contained a Gulliver-related transposable element in the 9^th^ exon of the *METE* gene, was completely reliant on B_12_ for growth but in the presence of the vitamin it was able to outcompete its B_12_-independent progenitor. In the absence of B_12_ the *METE* mutant would sometimes revert to B_12_ independence and resume growth. Reversion was found to be due to excision of the transposon to leave behind a wild-type *METE* gene sequence, but there was a single case where 9 bp fragment of the transposon was left behind resulting in a stable B_12_-dependent strain hereafter called metE7.

*C. reinhardtii* is a well-researched model organism that has been instrumental in improving our understanding of algal photosynthesis, ciliogenesis, and responses to fluctuating nutrient environments (18–20). We wanted to use the metE7 mutant of *C. reinhardtii* to study how recently acquired B_12_ auxotrophy impacts an organism’s fitness and physiology, and to provide insight into the metabolic challenges that other B_12_ dependent algae might have faced when they first evolved. In this work we characterized the responses of metE7 to different vitamin B_12_ regimes and compared them to the responses of its ancestral B_12_-independent strain, and to a closely related, naturally B_12_-dependent alga *Lobomonas rostrata*. The responses of metE7 to B_12_ deprivation were quantified by measuring changes in gene expression, cellular composition, photosynthetic activity and viability, and were contrasted against changes under nitrogen deprivation. To assess whether a recently evolved algal B_12_ auxotroph could improve its survival during B_12_ deprivation relatively quickly, we subjected metE7 to a further experimental evolution period of several months in limited B_12_ or coculture with a B_12_ producing bacterium and characterised the resulting lines.

## Materials and Methods

### Strains

*Mesorhizobium loti* (MAFF 303099) was a gift from Prof. Allan Downie at the John Innes Centre, Norwich, UK. Algal strains used in this study are shown in Table S1 and include *Lobomonas rostrata* (SAG 45/2), as well as several *Chlamydomonas reinhardtii* strains derived from strain 12 of wild type 137c or the cell wall-deficient strain cw15. The stable B_12_-dependent metE7, the unstable B_12_-dependent (S-type) as well as the B_12_-independent revertant line (R-type) all evolved from the strain 12 of wild type 137c (Ancestral) as described by Helliwell et al. (2015). Another B_12_-dependent mutant (metE4) was generated by targeted (CRISPR/Cpf1) knockout of the *METE* gene in the UVM4 strain using the protocol described in Ferenczi et al. 2017 (21),

### Culture conditions and growth measurements

Algal colonies were maintained quarterly on Tris-acetate phosphate (TAP) + 1000 ng·l^−1^ cyanocobalamin (B_12_) agar (1.5%) in sealed transparent plastic tubes at room temperature and ambient light. Cultures were grown in TAP or Tris min medium under continuous light or a light-dark period of 16hr-8hr, at 100 µE·m^−2^·s^−1^, at a temperature of 25°C, with rotational shaking at 120 rpm in an incubator (InforsHTMultitron; Switzerland). For nutrient starvation experiments the pre-culture TAP media contained 200 ng·l^−1^ of B_12_, and when cell densities surpassed 1*10^6^ cells·ml^−1^ or an OD730 nm of 0.2, cultures were centrifuged at 2,000 g for 2 minutes, followed by supernatant removal and resuspension of the cell pellet in media.

Algal cell density and optical density at 730 nm were measured using a Z2 particle count analyser (Beckman Coulter Ltd.) with limits of 2.974-9.001 µm, and a FluoStar Optima (BMG labtech) or Thermo Spectronic UV1 spectrophotometer (ThermoFisher) respectively. Mean cell diameter was also quantified on a Z2 particle analyser (Backman Coulter Ltd.). Dry mass was measured by filtering 20 ml of culture through pre-dried and weighed grade 5 whatmann filter paper (Sigma-Aldrich WHA1005090), drying at 70°C for 24 hours, followed by further weighing on a Secura mass balance (Sartorius). Algal and bacterial CFU·ml were determined by plating on solid media.

### Measurement of photosynthetic parameters

200 µl of cultures with an OD730 nm>0.1 were transferred to a 96 well plate which was then incubated at 25°C in the dark for 20 minutes. F_0_ was measured prior to, and F_m_ during, a saturating pulse at 6172 µE·m^−2^·s^−1^. The light intensity was increased to 100 µE·m^− 2^·s^−1^ and the cells allowed to acclimate for 30 seconds prior to another set of fluorescence measurements before and during a saturating pulse. From these fluorescence measurements the CF imager software calculated non-photochemical quenching (*Fm/Fm’-1*), PSII maximum efficiency (*Fv’/Fm’*), and the coefficient of photochemical quenching (*Fq’/Fv’*) at each light intensity.

### Measurement of cellular biochemical composition

Lipids were extracted from the cell pellet from 10 ml of culture using the chloroform/methanol/water method and triacylglycerides (TAGs), polar lipids and free fatty acids in the total lipid extract and total fatty acid methyl esters (FAMEs) were analysed by GC-FID and GC-MS, as described in Davey et al. (2014) (22). A 1 ml aliquot of algal culture was used for pigment and starch quantification as described in Davey et al. (2014), and a 10 ml aliquot for protein quantification using a Bradford assay and amino acids by HPLC as described in Helliwell et al. (2018) (23).

### Reactive oxygen species quantification

2 µl of 1 mM 2’,7’ Dichlorofluorescein diacetate (Sigma-Aldrich) dissolved in DMSO was added to 198 µl of cell culture in a black f-bottom 96 well plate (Greiner bio-one) and incubated at room temperature in the dark for 60 minutes before recording fluorescence at 520 nm after excitation at 485 nm in a FluoStar Optima Spectrophotometer (BMG labtech). Fresh cell culture media devoid of any cells was used as a blank.

### SAM and SAH quantification

10 ml of samples were centrifuged at 2,000 g for 2 minutes, supernatant removed, and cell pellet lyophilised at <-40°C and <10 pascals for 12-24 hours. 300 µl of 10% methanol (LC-MS grade) spiked with stable isotope-labelled amino acids (L-amino acid mix, Sigma-Aldrich, Co., St. Louis, MO, USA) was added to each sample. They were vortexed 3 times, every 10 min, before sonicating for 15 min in an iced water bath then centrifuging (16,100 x g) for 15 min at 4^°^C. Quantitative analysis was performed on 150 µl of supernatant using an Agilent 6420B triple quadrupole (QQQ) mass spectrometer (Agilent Technologies, Palo Alto, USA) coupled to a 1200 series Rapid Resolution HPLC system. Details of the HPLC-MS are given in the supplementary information.

### Transcript quantification

Total RNA extraction was performed on the cell pellet from 10 ml of algal culture using the RNeasy® Plant Mini Kit (QIAGEN). DNase treatment was carried out using TURBO DNA-free™ kit (Ambion), and cDNA synthesis using SuperScript®III First-Strand synthesis system for RT-PCR (Invitrogen) according to the manufacturer’s instructions. RT-qPCR was performed as described by Helliwell et al. 2018 (24), using primers listed in Table S2

### Artificial Evolution setup

A culture of metE7 cells was plated on TAP +1000 ng·l^−1^ B_12_ agar, then 8 colonies picked and resuspended in TAP + 200 ng·l^−1^ B_12_ in a 96 well plate. Each well was split into 3 wells, each in a different 96 well plate containing 200 µl of a different media: TAP +1000 ng·l^− 1^ B_12,_ TAP +25 ng·l^−1^ B_12_, and TP medium. *M. loti* was prepared in a similar manner to metE7, except preculturing was performed in TP + 0.01% glycerol. *M. loti* was added to the TP culture containing metE7 at a density roughly 20 times greater than the alga. The 96 well plates were incubated at 25°C, under continuous light at 100 µE·m^−2^·s^−1^, on a shaking platform at 120 rpm. Each week the cultures were diluted: Those in TAP +1000 ng·l^−1^ B_12_ were diluted 10,000-fold, TAP +25 ng·l^−1^ B_12_ = 100-fold, and TP = 5-fold. Every three weeks 10 µl of serial dilutions of each culture was also spotted onto TAP agar + Ampicillin (50 µg·ml^−1^) and Kasugamycin (75 µg·ml^−1^) and TAP agar + 1000 ng·l^−1^ B_12_ to check for B_12_-independent *C. reinhardtii*, or bacterial contaminants and to act as a reserve in the case of contamination. If cultures were found to be contaminated, then at the next transfer they were replaced by colonies from the same well that had grown on the TAP agar plates. At four points during the 12-month evolution period all cultures were transferred to TAP agar plates where they were stored for 2 weeks during an absence from the lab, meaning that the total time in liquid culture was 10 months. See Fig S9 for an illustration of the experimental evolution setup and the tests of B_12_ dose response and viability during B_12_ deprivation that were performed on the evolved lines.

## Results

### B_12_ deprivation causes substantial changes to C1 metabolism in the metE7 mutant

Methionine synthase plays a central role in the C1 cycle (Fig. 1A), and thus facilitates nucleotide synthesis and production of the universal methyl donor S-adenosylmethionine, which is essential for many biosynthetic and epigenetic processes (25, 26). Wild-type (WT) *C. reinhardtii* can operate these cycles in the absence of B_12_ using the methionine synthase variant METE, but metE7 relies solely on the B_12_-requiring METH isoform. Before investigating the effect of B_12_ deprivation on C1 metabolism in metE7 we first wanted to eliminate the possibility that other mutations in the experimentally evolved metE7 line might account for its B_12_ dependent phenotype. We therefore generated an independent *METE* mutant line (metE4) using CRISPR/Cpf1 (21). This mutant has an in-frame stop codon (Fig. S1) and, as predicted, exhibits B_12_-dependence. We therefore proceeded to investigate the effect of B_12_ on C1 metabolism in the metE7 line, since its origin is a closer reflection of how B_12_ auxotrophy would have arisen naturally in other algae.

**Figure 1.**
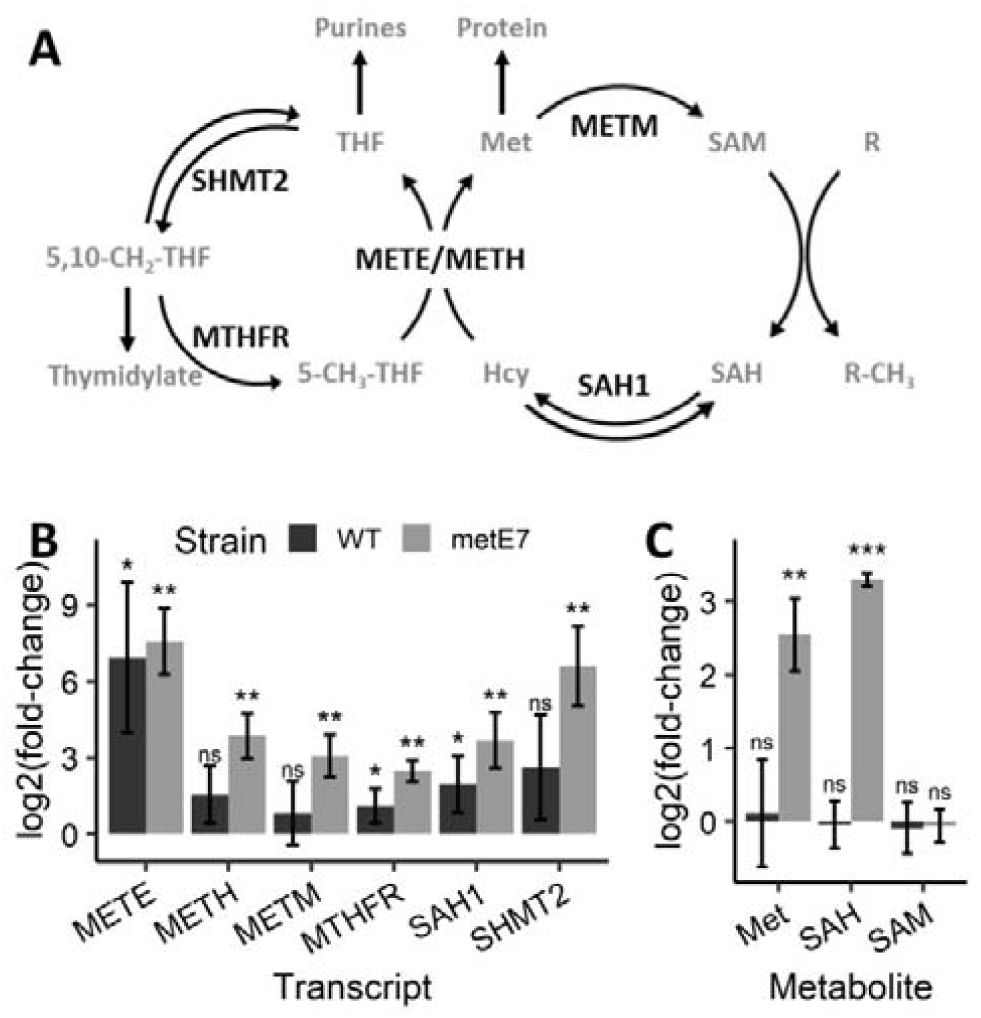
C1 cycle metabolites and transcripts increase during B_12_ deprivation of metE7. **(A)** Metabolic map of a portion of the C1 cycle centred around METE and METH, with enzyme abbreviations in black, metabolite abbreviations in grey, and arrows depicting enzyme-catalysed reactions. **(B)** Abundance of six transcripts for enzymes of the C1 cycle measured by RT-qPCR on RNA extracted from the ancestral line and metE7 after 30 hours of incubation in mixotrophic conditions with (1000 ng·I^−1^) or without B_12_. **(C)** Abundance of Met, SAM, and SAH metabolites measured by HPLC-MS on the same samples as above. Metabolite and transcript abundances are expressed as levels in B_12_-deprived conditions relative to B_12_-replete conditions and presented on a log_2_() scale. Error bars = sd, n=3-4, ‘ns’=not significant, *=p<0.05, **=p<0.01, ***=p<0.001, Welch’s t test. WT = ancestral B_12_-independent strain, *metE7* = experimentally evolved B_12_-dependent line. See also figure S2.

Both the WT ancestral line and metE7 were precultured in TAP medium in continuous light with adequate (200 ng·l^−1^) B_12_ to maintain a low cellular quota of the vitamin. The cells were then pelleted, washed and transferred to B_12_ replete (1000 ng·l^−1^) or B_12_ deprived (no B_12_) TAP medium at 5×10^5^ cells/ml and grown for 30 hours. Steady state transcript levels of six enzymes in the C1-cycle were then investigated by RT-qPCR (Fig. 1*B*). In the WT, three transcripts (*METE, SAH1*, and *MTHFR*) were significantly (p<0.05) upregulated by B_12_ deprivation, while in metE7 all six (including *METH, METM*, and *SHMT2*) increased. Levels of the methionine cycle metabolites methionine, SAM and SAH were quantified by HPLC-MS. In the WT there was no difference in methionine, SAM or SAH levels in the two conditions (Fig. 1C). However, in metE7 cells under B_12_ deprivation methionine levels were raised 6-fold, which was somewhat unexpected given that methionine synthase activity was impeded. SAH levels were also significantly elevated, whereas there was no effect on SAM. Consequently, the SAM:SAH ratio decreased by 10-fold to 3:1 under B_12_ deprivation. We then studied the dynamics of these changes by measuring metabolites and RNA abundance at several points during 3 days of B_12_ deprivation and then for 2 days following add-back of 1000 ng·l^−1^ B_12_. The transcripts for all six tested C1 cycle genes increased rapidly in the first 6 h and then plateaued; reintroduction of B_12_ led to an immediate reduction to near initial amounts (Fig. S2A). Similar profiles were seen for the metabolites SAM and SAH, although the peak occurred later at 24 h (Fig. S2B). Methionine levels were more variable, but nonetheless there was a similar trend of a peak 24 h after removal of B_12_. More significantly, the SAM:SAH ratio fell sharply from 30 to less than 1 within 24 h. A subsequent gradual increase occurred over the next 2 days, and resupply of B_12_ increased this ratio further over the following 2 days. The likelihood therefore is that many cellular processes would be impacted in B_12_-deprived metE7 cells.

### B_12_ deprivation significantly impacts cell physiology and biochemical composition

Our data demonstrate a substantial impact of B_12_ limitation on the expression of C1 metabolic genes as well as the abundance of C1 metabolites. To elucidate downstream consequences of perturbed C1 metabolism we also characterised broader physiological responses to B_12_ deprivation. As has been documented previously (17), growth of metE7 cells was significantly impaired in B_12_-deprived conditions (Fig. S3A). However, by day 2 the B_12_ deprived cells had a 36% larger diameter resulting in a 150% increase in volume (Fig. 2A and Fig. S3B), indicating that cell division was more restricted than overall growth. Moreover, cell viability, which was assayed by the ability of cells to form colonies when plated on B_12_-replete TAP agar, decreased to below 25% within 4 days of B_12_ limitation (Fig. S3C). This was preceded by a reduction in photosystem II maximum efficiency (*Fv/Fm*) (Fig. S3D), an often-used indicator of algal stress (27, 28).

**Figure 2.**
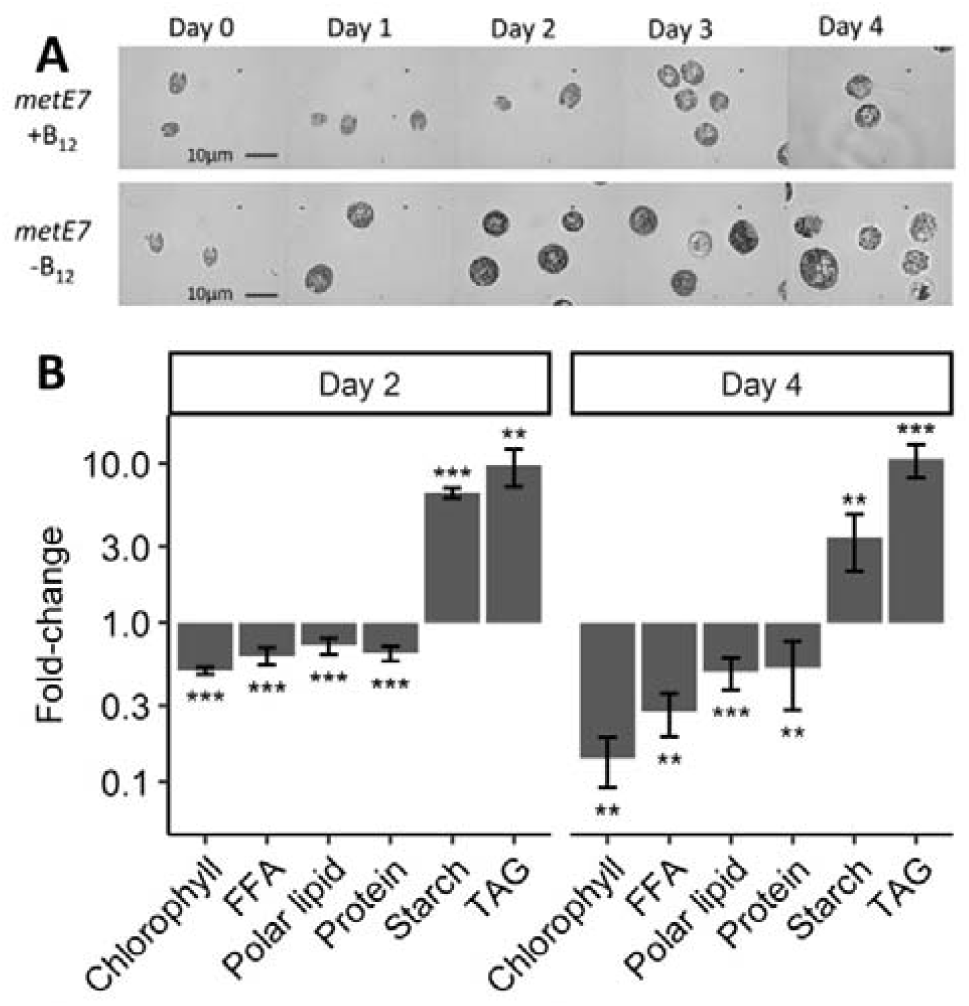
B_12_ deprivation of *metE7* causes cell enlargement and significant changes in macromolecular composition. **(A)** Microscope photographs taken at 1000× magnification of *metE7* cells grown in TAP medium in B_12_ replete (1000 ng·l^−1^) or B_12_ deprived (0 ng·l^−1^) conditions over a period of 4 days **(B)** Macromolecular composition of B_12_-deprived cells on day 2 and day 4 of the growth period expressed as mass of those compounds normalised to total cell dry mass and then expressed relative to the amounts in B_12_ replete conditions. Error bars = sd, n = 5. **=p<0.01, ***=p<0.001, Welch’s t test.

The biochemical composition of *C. reinhardtii* cells is altered considerably and similarly under various nutrient deprivations and so we hypothesised that B_12_ limitation would also induce broadly the same responses (20, 29, 30). Therefore, metE7 cells were precultured as before in 200 ng·l^−1^ B_12_, then washed and resuspended in TAP with (1000 ng·l^−1^) or without B_12_ and cultured mixotrophically for 4 days. Cultures were visually inspected by microscopy (Fig. 2*A*) and the amounts of various cellular components were measured on day 2 and 4 (Fig. 2*B*). Chlorophyll levels declined considerably under B_12_ deprivation so that by day four the cells had a bleached appearance with an 85% lower concentration than the B_12_ replete cells. Similarly, free fatty acids (FFA), polar lipids and proteins were at least 50% lower under B_12_ deprived conditions on day 4. Starch content on the other hand, showed the largest absolute increase from B_12_ replete to B_12_ deprived cells (Fig S3), and triacylglycerides were 10-fold higher in B_12_-deprived cells (Fig 2B), which effectively balanced the loss of polar lipids and free fatty acids so that overall lipid levels were roughly 8-10% of dry mass in both treatments. To look in more detail, quantification of free amino acids and fatty acid composition of all lipid classes was carried out (Fig S4). By day 4 most of the amino acids decreased significantly under B_12_ deprivation. Particularly noteworthy is the reduction in methionine, in contrast to its elevation at an earlier timepoint, and the increase in glutamine, the only amino acid to be more abundant in B_12_ deprived cells. Overall the degree of fatty acid saturation was higher under B_12_ deprivation, due mainly to an increase in the dominant saturated fatty acids palmitate (16:0) and stearate (18:0) (Fig. S5B), although levels of several unsaturated fatty acids, in particular 16:2, 16:3^(7,10,13)^, 18:1 and 18:2, were also elevated.

### Responses to nitrogen deprivation improve survival under B_12_ deprivation

Our results demonstrate that B_12_ deprivation of metE7 causes several changes in biochemical composition akin to those exhibited following nitrogen deprivation of WT *C. reinhardtii*. To further investigate this comparison we measured growth, viability, and photosynthetic efficiency under both conditions over a timecourse (Fig. S6). metE7 culture density increased more under B_12_ than nitrogen deprivation (Fig. S6A), but started to decline after day 2, unlike under nitrogen deprivation where growth continued more slowly over 4 days. For cell viability, both conditions caused a decline, but while loss of viability continued in B_12_ deprived cells, under nitrogen deprivation the initial loss was followed by recovery (Fig. S6B). Maximum photosynthetic efficiency of photosystem II, however, did not recover under either condition, and its decline was more rapid in nitrogen-deprived cells (Fig. S6C).

The increased viability of metE7 under nitrogen compared with B_12_ deprivation suggested to us that either the metabolic role of B_12_ would make it intrinsically more difficult to cope without or that the evolutionary naivety of metE7 to B_12_ dependence would mean it had little time to evolve protective responses to B_12_ limitation. We therefore tested whether responses to nitrogen deprivation could afford some protection against B_12_ deprivation. Viability measurements were monitored over several days, and cultures lacking nitrogen or B_12_ behaved as previously (Fig 3A). However, metE7 cells deprived of both nitrogen and B_12_ simultaneously were more similar to those starved on nitrogen: there was an initial decrease in viability followed by recovery to a level significantly higher than in B_12_ deprivation alone. As total growth in B_12_ and nitrogen deprivation was not significantly different from B_12_ deprivation alone (Fig. S7) this apparent protective mechanism in response to nitrogen deprivation is not simply a result of inhibiting growth and hence avoiding severe B_12_ starvation.

**Figure 3.**
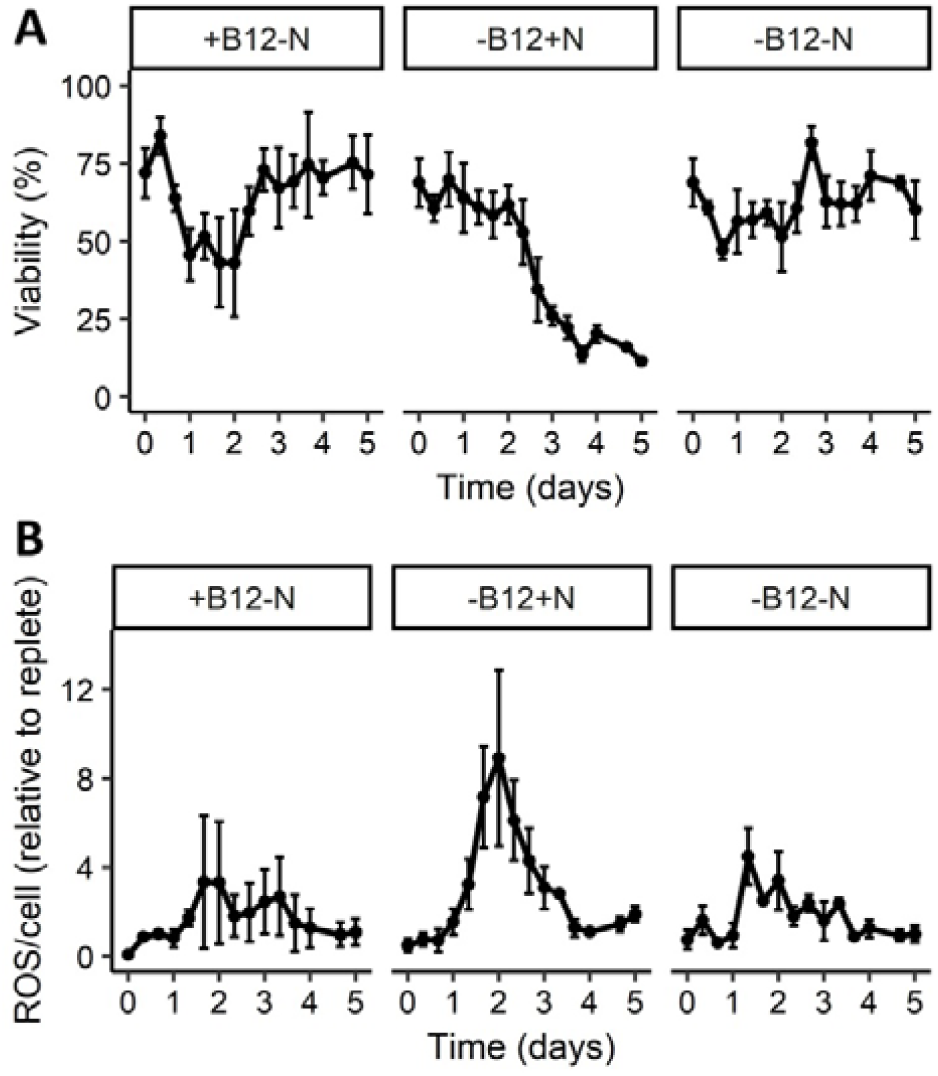
*metE7* survives better and produces lower levels of reactive oxygen species (ROS) when limited for both N and B_12_ than just B_12_ alone. **(A)** Percentage of cells that could form colonies (a measure of viability) on nutrient replete agar when removed at different timepoints from nutrient deprivation conditions (Indicated in panels above the graphs). **(B)** Reactive oxygen species (ROS) measured by dichlorofluorescein diacetate (DCFDA) fluorescence and normalised both on a per cell basis and to the nutrient replete treatment (+B_12_+N). Error bars = sd, n = 3-6.

In *C. reinhardtii*, as in many photosynthetic organisms, the absorption of light energy in excess of that required for metabolism can increase the production of reactive oxygen species (ROS) (31). To investigate whether the cell death observed under B_12_ deprivation of metE7 could be due to ROS, the general ROS-sensitive dye dihydrodichlorofluorescein diacetate was incubated with cells at different timepoints during nutrient deprivation. We found that ROS levels increased in all nutrient deprived conditions in the first two days but were highest in those cells deprived of B_12_ alone (Fig. 3B). This peak coincided with the start of the substantial decline in cell viability (Fig. 3A). The combination of B_12_ and nitrogen deprivation reduced ROS levels to similar amounts to those seen in the nitrogen-deprived cells, and so may be a factor behind reduced cell death.

### Natural B_12_ auxotroph Lobomonas rostrata fares better under B_12_ limiting conditions than metE7

Considering that metE7 quickly lost viability in the absence of B_12_ while nitrogen starvation invoked protective responses independent of B_12_ status, it is possible that as a novel auxotroph metE7’s response to B_12_ deprivation is simply underdeveloped. To test this we compared the B_12_ physiology of metE7 with *Lobomonas rostrata*, a naturally B_12_-dependent member of the same Volvocaceae family of chlorophyte algae (32, 33). Cell viability was significantly greater in *L. rostrata* cells compared to the metE7 line after 2-4 days of B_12_ deprivation despite also growing to a greater density (Fig S7A). Moreover, a B_12_ dose-response experiment, in which the two species were each cultured mixotrophically in a range of B_12_ concentrations, revealed that *L. rostrata* reached a higher optical density than metE7 at all B_12_ concentrations below 90 ng·l^−1^, while the inverse was true above 90 ng·l^−1^ (Fig 4A). This indicates that *L. rostrata* has a lower B_12_ requirement than metE7.

**Figure 4.**
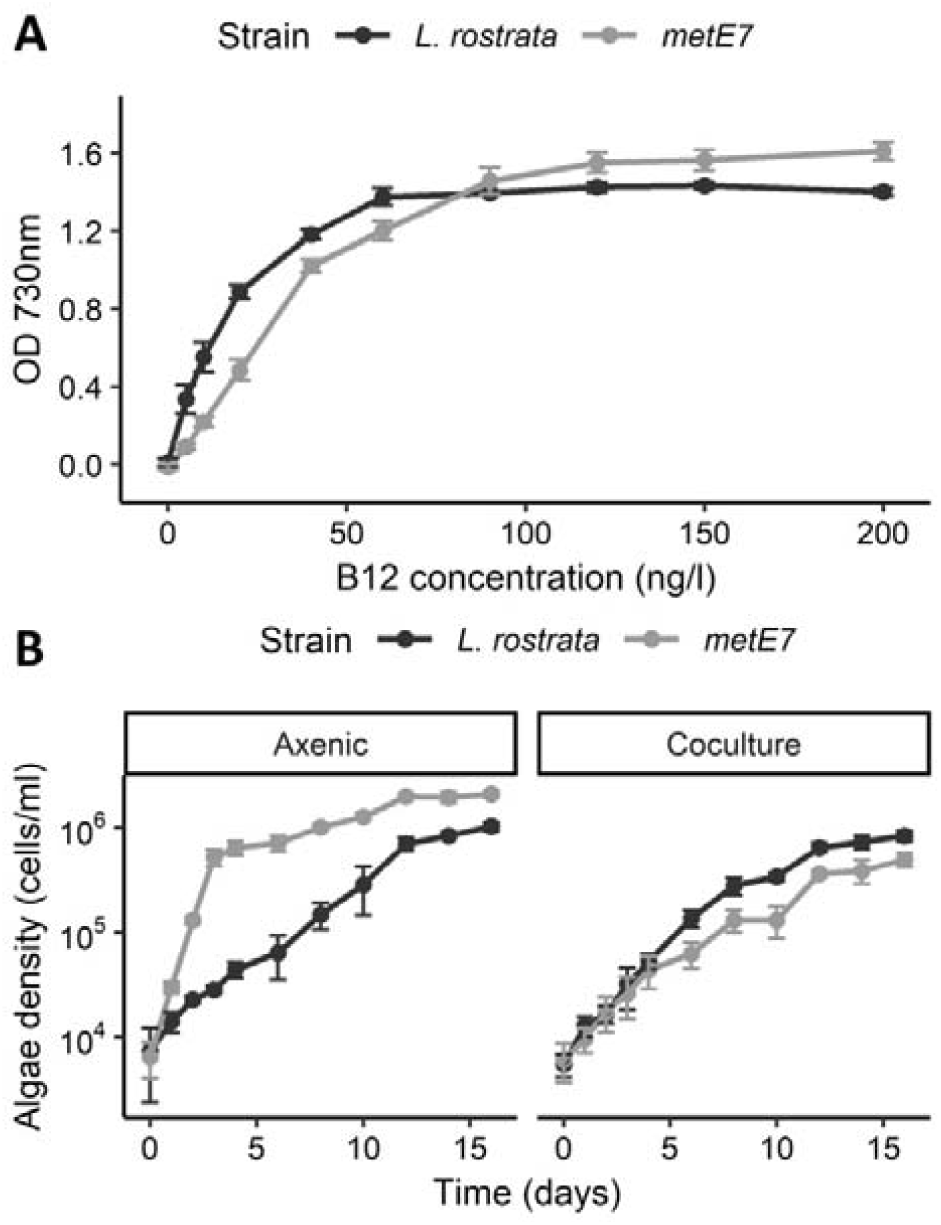
*L. rostrata* grows better than *metE7* in coculture with a B_12_ producing bacterium, in part due to its lower demand for B_12_. **(A)** Cultures were grown mixotrophically (TAP medium in continuous light), B_12_ concentrations ranged from 0 to 200 ng·l^−1^ and precultures of the algae, which were grown with 200 ng·l^−1^ B_12_, were washed thrice and inoculated at a density of roughly 100 cells·ml^−1^. Culture density was measured as optical density at 730 nm after 5 days of growth for the *C. reinhardtii* strains and 9 days for *L. rostrata*. **(B)** Cultures were grown photoautotrophically (Tris minimal media in 16h:8h light:dark cycles) in axenic culture (with 100 ng·l^−1^ B_12_) or coculture (with the B_12_-producing bacterium *M. loti*) over a period of 16 days with measurements of cell density every 1-2 days. For both panel A and B, black = *L. rostrata*, grey = *metE7*, error bars = sd, n=4.

In the natural environment the ultimate source of B_12_ is from prokaryotes since they are the only known B_12_ producers (34). In separate studies it was shown that B_12_-dependent growth of *L. rostrata* and metE7 can be supported by the B_12_ synthesising bacterium *Mesorhizobium loti* (17, 35). We therefore compared directly the growth of metE7 and *L. rostrata* in B_12_-supplemented (100 ng·l^−1^) axenic culture and in coculture with *M. loti* in media lacking a carbon source (TP) (Fig 4A). Even though metE7 grew much more quickly and to a higher density than *L. rostrata* under axenic, B_12_-supplemented conditions, it grew less well in coculture with *M. loti* (Fig. 4B), indicating B_12_ provision from the bacterium is less effective at supporting the growth of metE7 than of *L. rostrata*, perhaps simply due to their different B_12_ requirements, but possibly due to more sophisticated symbiotic interactions.

### Experimental evolution in coculture improves B_12_-use efficiency and resilience to B_12_ deprivation

Together our data suggest that the newly evolved metE7 line is poorly adapted to coping with B_12_ deprivation, but we wanted to determine whether the metE7 line could evolve improved tolerance to B_12_ limiting conditions, so we employed an experimental evolution approach. We designed three distinct conditions, referred to as H, L and C. Condition H (TAP medium with high (1000 ng·l^−1^) B_12_) was a continuation of the conditions that had initially generated *metE7* (17). Condition L (TAP medium with low (25 ng·l^−1^) B_12_) was chosen so that B_12_ would limit growth. Condition C (coculture with *M. loti* in TP medium) was a simplification of an environmental microbial community. Eight independent cultures for each condition were established from a single colony and then subcultured once per week over a total period of 10 months. To account for the different growth rates in the three conditions, we applied the following dilution rates of 10,000, 100, and 5 times per week in condition H, L and C respectively (Fig S8). After 10 months under selective conditions all 24 cultures had survived and were then treated with antibiotics to remove the *M. loti* from condition C and to ensure that there were no other contaminating bacteria. We then subcultured the lines in mixotrophic conditions with TAP + 200 ng·l^−1^ B_12_ three times over nine days to ensure they were all acclimated to the same conditions. The behaviours of the algal populations, hereafter referred to as metE7H, metE7L, and metE7C, were then compared alongside the progenitor metE7 line, which had been maintained on TP agar with 1000 ng·l^−1^ B_12_ without subculturing.

Under high levels of B_12_ (320 ng·l^−1^) a similar optical density was reached by the progenitor metE7 strain and the metE7H and metE7C populations, whereas metE7L growth was somewhat compromised (Fig. S10A). When grown across a range of B_12_ concentrations to determine a dose response, the metE7C populations reached a significantly higher optical density at the lower concentrations of 20 and 40 ng·l^−1^ B_12_ than the other lines(Fig. 5A). The concentration of B_12_ required to produce half the maximum growth (EC_50_) of metE7C was therefore much lower than the progenitor metE7 or metE7H (Fig. S10B) and this was reflected in the higher B_12_ use efficiency i.e. the maximal increase in yield (OD_730_) that results from an increase in B_12_ concentration (Fig. 5B). However, the maximal growth rate of metE7C was significantly lower (Fig. S10C), and it is tempting to conclude that this is a necessary trade-off. We also compared the viability of the experimentally evolved lines during B_12_ deprivation (Fig. 5C). Fig. 5C shows that although all lines lost viability during B_12_ deprivation, metE7L and metE7C survived substantially better, with a median survival time more than a day longer (Fig. 5D) than both the progenitor metE7 and metE7H.

**Figure 5.**
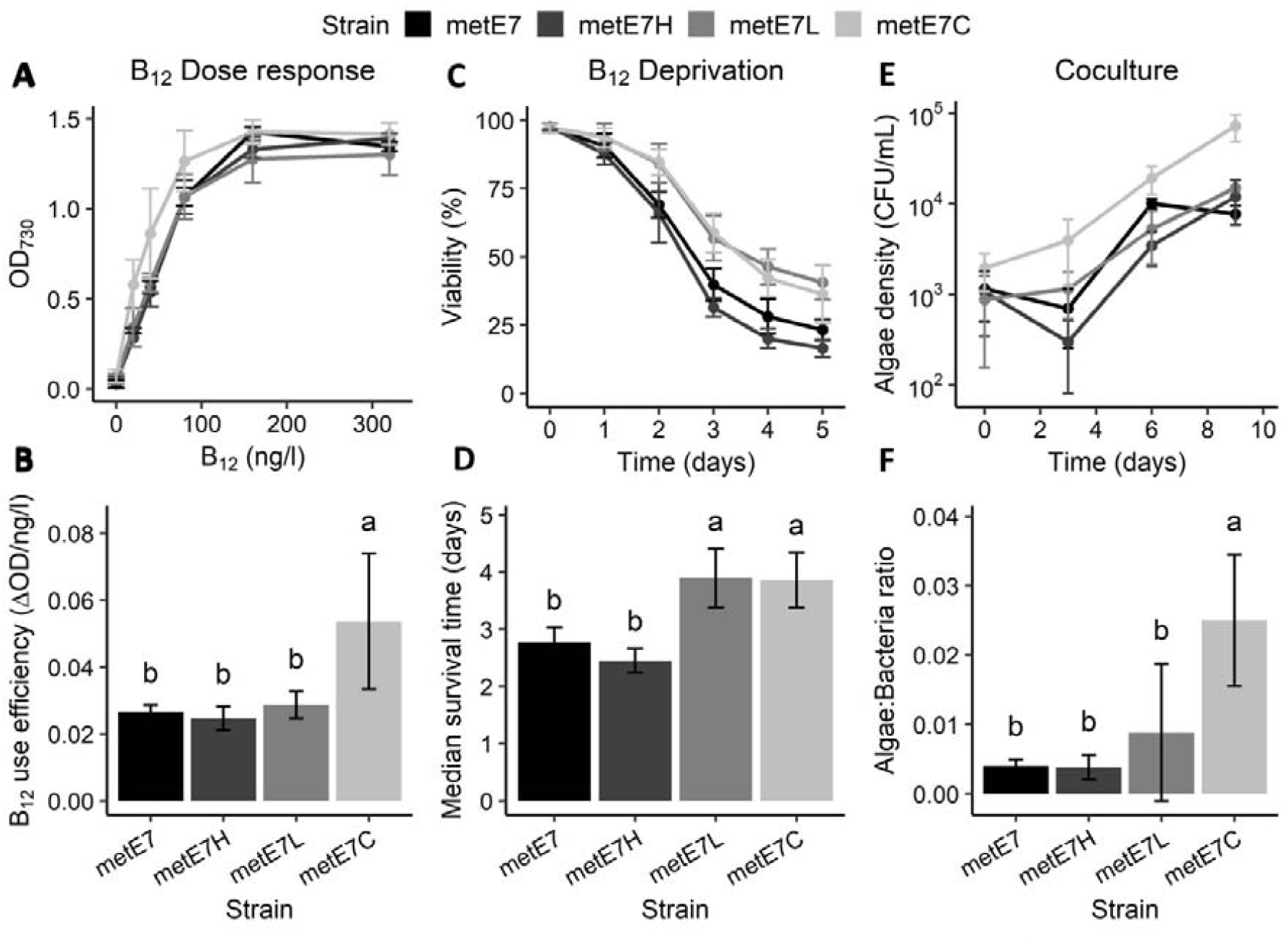
Experimental coevolution of metE7 with the bacterium *M. loti* selects for improved algal growth in coculture, increased B_12_ use efficiency and better resilience to B_12_ deprivation. **(A)** Maximum optical density achieved by mixotrophically-grown cultures of experimentally evolved lines of metE7 grown over a period of 12 days in six different concentrations of B_12_. **(B)** B_12_ use efficiency of evolved lines calculated using a fitted Monod equation and expressed as the maximum rate of increase in OD_730_ that would result from an increase in B_12_ concentration. **(C)** Viability (measured as the percentage of cells capable of forming colonies on B_12_-replete agar) of mixotrophically-grown cultures of experimentally evolved lines of metE7 over a 5-day period cultured in 40 ng· l^−1^ B_12_. **(D)** Median survival time of evolved lines after dilution of culture to 40 ng· l^−1^ B_12_ calculated using a fitted Verhulst equation. **(E)** Algal cell density of photoautotrophically-grown cocultures of experimentally evolved lines of metE7 with *M. loti* over a 9-day period. **(F)** Ratio of algae to bacteria on the final day (day 9) of growth in coculture. metE7H = metE7 evolved in TAP + 1000 ng· l^−1^ B_12_ for 10 months, metE7L = metE7 evolved in TAP media + 25 ng· l^−1^ B_12_, metE7C = metE7 evolved in Tris minimal medium in coculture with the B_12_-producing bacterium *M. loti*. Error bars = 95% confidence interval, n = 7-8, letters above error bars indicate statistical groupings provided by Tukey’s test, which was performed following a significant ANOVA result.

To elucidate which factors contributed to the improved survival, we performed a multi-parameter physiological analysis (Fig. S11). 16 parameters were measured across the 32 metE7 populations and the dataset visualised in three ways. Fig S11A presents the data as a heatmap with the most similar populations, which generally were those exposed to the same evolution conditions, clustered together to form a phylogenetic tree. Fig. S11B displays the first two components of a principal component analysis of the data, which confirmed that the experimental evolution populations tended to form separate clusters. Fig. S11C is a correlation matrix of the parameters to reveal those pairs that are most positively or negatively correlated with one another. A more definitive statistical approach was then used to determine the most important parameters for predicting survival time during B_12_ deprivation: Using stepwise minimisation of the Bayesian information criterion of the full linear model the 15 other parameters were reduced to just three. So, it was concluded that higher B_12_ use efficiency, lower ROS levels and lower maximal growth rate were sufficient to explain longer survival time under B_12_ deprivation of the metE7 populations.

Comparison of the growth of the evolved lines when cocultured with *M. loti* showed, perhaps unsurprisingly, that the metE7C lines grew better than the others (Fig 5E), and at the end of the growth period had a significantly higher number of algae supported per bacterium (Fig 5F). This algal:bacterial ratio was also optimally predicted by three parameters: higher algal B_12_ use efficiency and lower algal maximal growth rate, as for survival time, but also lower algal B_12_ uptake capacity. Together these results indicate that experimental evolution in coculture not only improves growth in coculture but also increases B_12_ use-efficiency and survival during B_12_ deprivation.

## Discussion

In this study we exploited a novel model system for the evolution of vitamin B_12_ dependence by analysing the physiological and metabolic responses to B_12_ deprivation of an artificially evolved B_12_-dependent mutant of *C. reinhardtii*. Our analyses demonstrate that B_12_ deprivation has important consequences for C1 metabolism: we observed a significant increase in the transcript abundance of C1 cycle enzymes in both the wild type and metE7 strain, and a decrease in the methylation index (SAM:SAH ratio) in metE7 only. Moreover, B_12_ deprivation of metE7 causes a decrease in chlorophyll, protein and amino acids, and an increase in starch, lipids and saturated fatty acids, characteristic of limitation responses to macronutrients such as nitrogen. The rapid loss of viability seen under B_12_ deprivation can be averted if the metE7 cells are also limited for nitrogen, suggesting that it is not the lack of B_12_ *per se* that causes cell death, but an inability to respond appropriately. Together this suggests a newly evolved B_12_ auxotroph would be poorly adapted to surviving in the natural environment where a B_12_ supply is not guaranteed. However, we found that metE7 can be supported for several months by a B_12_-producing bacterium, and experimental evolution under these conditions caused improved B_12_ use efficiency and resilience to B_12_ deprivation.

B_12_ deprivation of metE7 decreased the SAM:SAH ratio 10-fold, similar to what was reported in a recent B_12_ deprivation study of the diatom *T. pseudonana* (13). As SAH is a competitive inhibitor of methyltransferases(36), this decrease would likely lead to general hypomethylation in metE7. The epigenetic marks methyldeoxyadenosine and methylcytosine are similarly abundant in *C. reinhardtii* and appear to mark active genes and repeat-rich regions respectively, so the consequences of hypomethylation are unclear (37, 38). The reduced abundance of B_12_-bound METH under B_12_ deprivation would hinder methionine synthesis and could cause the observed reduction in protein abundance (Fig. 2B). However, methionine levels increased between 12 and 24h of B_12_ deprivation (Fig S1B), suggesting a reduction in its use, proteolysis, or increased synthesis due to higher METH expression or via alternative pathways such as the S-methylmethionine cycle, as documented in plants (39).

*METE* transcript abundance showed a much higher dynamic range than *METH* during B_12_ deprivation and add-back (Fig. S2A), which is reflected by the higher diurnal range of *METE* observed in global transcriptomics and proteomics datasets (40). However, on average METE is around 60-fold more abundant than METH in *C. reinhardtii* (40). This may be due to a lower maximal catalytic rate of METE, as has been observed in *E. coli* (41), or due to its role in the flagella, which contain METE but not METH (42). Under B_12_ deprivation conditions the activity of METH would be compromised, yet in both metE7 and the ancestral strains it was upregulated. This is more similar to the B_12_ dependent algae *T. pseudonana* and *Tisochrysis lutea*, which also upregulate *METH* on B_12_ deprivation (12, 43), than the B_12_ independent *P. tricornutum*, which decreases *METH* expression (44). However, in both *T. pseudonana* and *P. tricornutum* B_12_ deprivation substantially upregulates C1 cycle enzymes including homologs of *METM, MTHFR* and *SAH1* (12), reflecting our findings and those of Helliwell et al. (2014) (45). Under sulfur and nitrogen deprivation conditions these C1 cycle genes are downregulated, suggesting that their upregulation during B_12_ deprivation is not a general response to nutrient stress, but a nutrient-specific one, as indeed is the case for *T. lutea* (43, 46, 47).

Chlorosis is a common symptom of nutrient deficiency in *C. reinhardtii*, evident in nitrogen, sulfur, iron, and zinc limiting conditions and so it is not surprising that B_12_ deprivation of metE7 caused a substantial decline in total chlorophyll (Fig. 2*B*) (48–50). The decrease in total protein content occurred more slowly and was less substantial (50% reduction over four days) than reported under nitrogen and sulphur deprivation (80% reduction within one day) (51). During nitrogen and iron starvation in *C. reinhardtii* membrane lipids decrease drastically concomitant with the increase in TAGs (52, 53). This is very much like what we observed for metE7 under B_12_ deprivation, although here the level of free fatty acids and polar lipids decreased by a roughly similar amount to the increase in TAGs indicating there is little to no *de novo* fatty acid synthesis. In addition, B_12_ deprivation causes similar shifts in fatty acid composition to nitrogen and iron deprivation, most notably a substantial increase in palmitic acid (16:0) and decrease in polyunsaturated 16:4 fatty acid (53, 54). Despite these similarities, B_12_ deprivation may elicit an increase in TAGs by a different pathway due to disrupted C1 metabolism, as has been observed in several organisms (55–57). This is thought to be due to a reduction in the methylation potential limiting membrane lipid synthesis and hence diverting more lipids towards TAGs (57, 58). Therefore, B_12_ deprivation could provide a complementary approach to other nutrient deprivation experiments in improving our understanding of lipid metabolism in *C. reinhardtii* and other algae.

From an evolutionary perspective, the prevalence of vitamin B_12_ dependence among algae appears somewhat at odds with the severe fitness penalties that would be incurred given limiting dissolved B_12_ concentrations, particularly when the fitness benefit in replete B_12_ is marginal (17). However, relative to optimal axenic laboratory conditions in which the metE7 line evolved, in the environment multiple nutrients may colimit growth perhaps even eliciting responses that mitigate against B_12_ deprivation, as we observed here, and B_12_-producing bacteria may not simply co-occur with algae but also actively engage in mutualistic interactions (1, 35, 59, 60). Furthermore, our evidence suggests that selection under coculture conditions led to the newly evolved B_12_ auxotroph developing increased B_12_ use efficiency and becoming better adapted to tolerating B_12_ limitation, which could make this line more robust to the unreliable B_12_ supply in the natural environment. However, these improvements appeared to come at the expense of maximal growth rate in B_12_ replete conditions (Fig. S10C), which is not unexpected in light of previous experimental evolution studies in *C. reinhardtii* (61). As one of the conserved responses of *C. reinhardtii* upon detecting depletion of various nutrients is to decrease cell division, it is possible that slower growth might even be selected for under B_12_ deprivation. Indeed, a low growth rate was found to be a significant predictor of greater survival time under B_12_ deprivation, alongside low ROS levels and high B_12_ use efficiency.

The fact that metE7 survived a 10-month period either with limited artificial supplementation of B_12_ or by relying completely on bacterial B_12_ provision, does suggest that even a newly evolved and poorly adapted B_12_ auxotroph would have ample opportunity to adapt further. What adaptations are likely to improve growth and survival under B_12_ deprivation are not altogether clear, but it is not unreasonable to assume that exaptation of existing nutrient limitation responses would play a major role. B_12_ dependence is certainly a risky evolutionary strategy, and one which may have ended in extinction countless times, but our work suggests that even the simplest of symbioses with B_12_-producing bacteria may be sufficient to ensure the survival and drive the continued evolution of B_12_-dependent algae.

## Acknowledgements

This work was supported by the BBSRC Doctoral Training Partnership BB/M011194/1 (FB, AGS), BBSRC BB/M018180/1 (PM, AGS), BB/I013164/1 (KEH & AGS), Leverhulme Trust RPG 2017-077 (MPD, AGS), Natural Environment Research Council NE/R015449/1 (KEH).

## Author contributions

F.B., P.M. and A.G.S designed the research; F.B., D.L.S. and A.H. performed the research; N.S., D.L.S. and M.P.D. contributed new reagents or analytic tools; F.B. and D.L.S. analysed the data; F.B., A.G.S., K.E.H. and P.M. wrote the paper with input from all authors.

## Parsed Citations

1. Croft MT, Lawrence AD, Raux-Deery E, Warren MJ, Smith AG (2005) Algae acquire vitamin B12 through a symbiotic relationship with bacteria. Nature 438(7064):90–3.

2. Panzeca C, et al. (2009) Distributions of dissolved vitamin B12 and Co in coastal and open-ocean environments. Estuar Coast Shelf Sci 85(2):223–230.

3. Sanudo-Wilhelmy SA, et al. (2012) Multiple B-vitamin depletion in large areas of the coastal ocean. Proc Natl Acad Sci 109(35):14041–14045.

4. Gobler C, Norman C, Panzeca C, Taylor G, Sañudo-Wilhelmy S (2007) Effect of B-vitamins (B1, B12) and inorganic nutrients on algal bloom dynamics in a coastal ecosystem. Aquat Microb Ecol 49:181–194.

5. Panzeca C, et al. (2008) Potential cobalt limitation of vitamin B12 synthesis in the North Atlantic Ocean. Global Biogeochem Cycles 22(2):n/a-n/a.

6. Ohwada K (1973) Seasonal Cycles of Vitamin B12, Thiamine and Biotin in Lake Sagami. Patterns of Their Distribution and Ecological Significance. Int Rev der gesamten Hydrobiol und Hydrogr 58(6):851–871.

7. Sañudo-Wilhelmy SA, Gobler CJ, Okbamichael M, Taylor GT (2006) Regulation of phytoplankton dynamics by vitamin B 12. Geophys Res Lett 33(4):L04604.

8. Bertrand EM, et al. (2007) Vitamin B12 and iron colimitation of phytoplankton growth in the Ross Sea. Limnol Oceanogr 52(3):1079–1093.

9. Browning TJ, et al. (2017) Nutrient co-limitation at the boundary of an oceanic gyre. Nature 551(7679):242.

10. Cohen NR, et al. (2017) Iron and vitamin interactions in marine diatom isolates and natural assemblages of the Northeast Pacific Ocean. Limnol Oceanogr 62(5):2076–2096.

11. Helliwell KE, Wheeler GL, Leptos KC, Goldstein RE, Smith AG (2011) Insights into the evolution of vitamin B12 auxotrophy from sequenced algal genomes. Mol Biol Evol 28(10):2921–33.

12. Bertrand EM, et al. (2012) Influence of cobalamin scarcity on diatom molecular physiology and identification of a cobalamin acquisition protein. Proc Natl Acad Sci 109(26):E1762–71.

13. Heal KR, Kellogg NA, Carlson LT, Lionheart RM, Ingalls AE (2019) Metabolic Consequences of Cobalamin Scarcity in the Diatom Thalassiosira pseudonana as Revealed Through Metabolomics. Protist 170(3):328–348.

14. Ellis KA, Cohen NR, Moreno C, Marchetti A (2017) Cobalamin-independent Methionine Synthase Distribution and Influence on Vitamin B12 Growth Requirements in Marine Diatoms. Protist 168(1):32–47.

15. Helliwell KE, Wheeler GL, Smith AG (2013) Widespread decay of vitamin-related pathways: coincidence or consequence? Trends Genet 29(8):469–78.

16. Helliwell KE (2017) The roles of B vitamins in phytoplankton nutrition: new perspectives and prospects. New Phytol 216(1):62–68.

17. Helliwell KE, et al. (2015) Fundamental shift in vitamin B12 eco-physiology of a model alga demonstrated by experimental evolution. ISME J 12:1–10.

18. Dubini A, Mus F, Seibert M, Grossman AR, Posewitz MC (2009) Flexibility in anaerobic metabolism as revealed in a mutant of chlamydomonas reinhardtii lacking hydrogenase activity. J Biol Chem 284(11):7201–7213.

19. Rochaix J-D (1995) Chlamydomonas reinhardtii as the Photosynthetic Yeast. Annu Rev Genet 29(1):209–230.

20. Grossman A (2000) Acclimation of Chamydomonas reinhardtii to its Nutrient environment. Protist 151(3):201–224.

21. Ferenczi A, Pyott DE, Xipnitou A, Molnar A Efficient targeted DNA editing and replacement in Chlamydomonas reinhardtii using Cpf1 ribonucleoproteins and single-stranded DNA. doi:10.1073/pnas.1710597114.

22. Davey MP, et al. (2014) Triacylglyceride production and autophagous responses in Chlamydomonas reinhardtii depend on resource allocation and carbon source. Eukaryot Cell 13(3):392–400.

23. Helliwell KE, et al. (2018) Quantitative proteomics of a B12-dependent alga grown in coculture with bacteria reveals metabolic tradeoffs required for mutualism. New Phytol 217(2):599–612.

24. Helliwell KE, et al. (2018) Quantitative proteomics of a B 12 -dependent alga grown in coculture with bacteria reveals metabolic tradeoffs required for mutualism. New Phytol 217(2):599–612.

25. Ducker GS, Rabinowitz JD (2017) Cell Metabolism Review One-Carbon Metabolism in Health and Disease. Cell Metab 25:27–42.

26. Lieber CS, Packer L (2002) S-Adenosylmethionine: molecular, biological, and clinical aspects-an introduction. Am J Clin Nutr 76(5):1148S–1150S.

27. White S, Anandraj A, Bux F (2011) PAM fluorometry as a tool to assess microalgal nutrient stress and monitor cellular neutral lipids. Bioresour Technol 102(2):1675–1682.

28. Parkhill J-P, Maillet G, Cullen JJ (2001) Fluorescence-based maximal quantum yield for PSII as a diagnostic of nutrient stress. J Phycol 37(4):517–529.

29. Saroussi S, Sanz-Luque E, Kim RG, Grossman AR (2017) Nutrient scavenging and energy management: acclimation responses in nitrogen and sulfur deprived Chlamydomonas. Curr Opin Plant Biol 39:114–122.

30. Juergens MT, Disbrow B, Shachar-Hill Y (2016) The Relationship of Triacylglycerol and Starch Accumulation to Carbon and Energy Flows during Nutrient Deprivation in Chlamydomonas reinhardtii. Plant Physiol 171(4):2445–57.

31. Erickson E, Wakao S, Niyogi KK (2015) Light stress and photoprotection in Chlamydomonas reinhardtii. Plant J 82(3):449–465.

32. Provasoli L (1958) Nutrition and Ecology of Protozoa and Algae. Annu Rev Microbiol 12(1):279–308.

33. Sausen N, Malavasi V, Melkonian M (2018) Molecular phylogeny, systematics, and revision of the type species of Lobomonas, L. francei (Volvocales, Chlorophyta) and closely related taxa. J Phycol 54(2):198–214.

34. Warren MJ, Raux E, Schubert HL, Escalante-Semerena JC (2002) The biosynthesis of adenosylcobalamin (vitamin B12). Nat Prod Rep 19(4):390–412.

35. Kazamia E, et al. (2012) Mutualistic interactions between vitamin B12 -dependent algae and heterotrophic bacteria exhibit regulation. Environ Microbiol 14(6):1466–76.

36. Chiang PK, et al. (1996) S-Adenosylmethionine and methylation. FASEB 10:471–480.

37. Fu Y, et al. (2015) N6-methyldeoxyadenosine marks active transcription start sites in Chlamydomonas. Cell 161(4):879–892.

38. Lopez D, et al. (2015) Dynamic Changes in the Transcriptome and Methylome of Chlamydomonas reinhardtii throughout Its Life Cycle. Plant Physiol 169(4):2730–43.

39. Ranocha P, et al. (2001) The S-methylmethionine cycle in angiosperms: ubiquity, antiquity and activity. Plant J 25(5):575–584.

40. Strenkert D, et al. (2019) Multiomics resolution of molecular events during a day in the life of Chlamydomonas. Proc Natl Acad Sci:201815238.

41. Gonzalez JC, Banerjee R V, Huang S, Sumner JS, Matthews RG (1992) Comparison of Cobalamin-Independent and Cobalamin-Dependent Methionine Synthases from Escherichia coli. Biochemistry (31):6045–6056.

42. Schneider MJ, Ulland M, Sloboda RD (2008) A Protein Methylation Pathway in Chlamydomonas Flagella Is Active during Flagellar Resorption. Mol Biol Cell 19(10):4319–4327.

43. Nef C, et al. (2019) How haptophytes microalgae mitigate vitamin B12 limitation. Sci Rep 9(1):8417.

44. Bertrand EM, et al. (2013) Methionine synthase interreplacement in diatom cultures and communities: Implications for the persistence of B 12 use by eukaryotic phytoplankton. 58(4):1431–1450.

45. Helliwell KE, et al. (2014) Unraveling Vitamin B12-Responsive Gene Regulation in Algae. Plant Physiol 165(1):388–397.

46. González-Ballester D, et al. (2010) RNA-Seq Analysis of Sulfur-Deprived Chlamydomonas Cells Reveals Aspects of Acclimation Critical for Cell Survival. Plant Cell 22(6):2058–2084.

47. Wase N, Black PN, Stanley BA, Dirusso CC (2014) Integrated Quantitative Analysis of Nitrogen Stress Response in Chlamydomonas reinhardtii Using Metabolite and Protein Profiling. J Proteome Res 13:1373–1396.

48. Schmollinger S, et al. (2014) Nitrogen-Sparing Mechanisms in Chlamydomonas Affect the Transcriptome, the Proteome, and Photosynthetic Metabolism. Plant Cell 26(4):1410–1435.

49. Juergens MT, et al. (2015) The regulation of photosynthetic structure and function during nitrogen deprivation in Chlamydomonas reinhardtii. Plant Physiol 167(2):558–73.

50. Kropat J, et al. (2011) A revised mineral nutrient supplement increases biomass and growth rate in Chlamydomonas reinhardtii. Plant J 66(5):770–780.

51. Cakmak T, et al. (2012) Differential effects of nitrogen and sulfur deprivation on growth and biodiesel feedstock production of Chlamydomonas reinhardtii. Biotechnol Bioeng 109(8):1947–1957.

52. Siaut M, et al. (2011) Oil accumulation in the model green alga Chlamydomonas reinhardtii: Characterization, variability between common laboratory strains and relationship with starch reserves. BMC Biotechnol 11(1):7.

53. Urzica EI, et al. (2013) Remodeling of membrane lipids in iron-starved chlamydomonas. J Biol Chem 288(42):30246–30258.

54. Msanne J, et al. (2012) Metabolic and gene expression changes triggered by nitrogen deprivation in the photoautotrophically grown microalgae Chlamydomonas reinhardtii and Coccomyxa sp. C-169. Phytochemistry 75:50–59.

55. Meï CE, et al. (2016) C1 Metabolism Inhibition and Nitrogen Deprivation Trigger Triacylglycerol Accumulation in Arabidopsis thaliana Cell Cultures and Highlight a Role of NPC in Phosphatidylcholine-to-Triacylglycerol Pathway. Front Plant Sci 7:1–16.

56. da Silva RP, Kelly KB, Al Rajabi A, Jacobs RL (2014) Novel insights on interactions between folate and lipid metabolism. BioFactors 40(3):277–283.

57. Visram M, et al. (2018) Homocysteine regulates fatty acid and lipid metabolism in yeast. J Biol Chem 293(15):5544–5555.

58. Malanovic N, et al. (2008) S-adenosyl-L-homocysteine hydrolase, key enzyme of methylation metabolism, regulates phosphatidylcholine synthesis and triacylglycerol homeostasis in yeast: Implications for homocysteine as a risk factor of atherosclerosis. J Biol Chem 283(35):23989–23999.

59. Cooper MB, et al. (2019) Cross-exchange of B-vitamins underpins a mutualistic interaction between Ostreococcus tauri and Dinoroseobacter shibae. ISME J 13(2):334–345.

60. Kazamia E, Helliwell KE, Purton S, Smith AG (2016) How mutualisms arise in phytoplankton communities: building eco-evolutionary principles for aquatic microbes. Ecol Lett 19(7):810–822.

61. Collins S, Bell G (2004) Phenotypic consequences of 1,000 generations of selection at elevated CO2 in a green alga. Nature 431(7008):566–569.

